# RF4Del: A Random Forest approach for accurate deletion detection

**DOI:** 10.1101/2022.03.10.483419

**Authors:** Roberto Xavier, Anna-Sophie Fiston-Lavier, Ronnie C.O. Alves, Emira Cherif

## Abstract

Efficiently detecting genomic structural variants (SVs) is a key step to grasp the “missing heritability” underlying complex traits involved in major evolutionary processes such as speciation, phenotypic plasticity, and adaptive responses. Yet, the SV-based genotype/trait association studies are still largely overlooked mainly due to the lack of reliable detection methods. Here, we present a random forest (RF) method for accurate deletion identification: RF4Del. By relying on the analysis of the mapping profiles, data already available in most sequencing projects, RF4Del can easily and quickly call deletions.

Several classic and ensemble learning strategies were carefully evaluated using proper benchmark data. RF4Del was trained and tested on simulated data from the model species Drosophila melanogaster to detect deletions. The model consists of 13 features extracted from a mapping file. We show that RF4Del outperforms established SV callers (DELLY, Pindel) with higher overall performance (F1-score > 0.75; 6x-12x sequence coverage) and is less affected by low sequencing coverage and deletion size variations. RF4Del could learn from a compilation of sequence patterns linked to a given SV. Such models can then be combined to form a learning system able to detect all types of SVs in a given genome, beyond the one used in our study. https://github.com/alvesrcoo/eletric-scheep

## Introduction

Genomic Structural Variants (SVs) are the major part of the variability between living organisms. SVs are three times more likely to be involved in genotype/complex trait association than a Simple Nucleotide Polymorphism (SNP), and SVs larger than 20 Kb are up to 50 times more likely to affect gene expression [1]. Indels account for 31% of all human genome mutations with deletions being twice the number of insertions [2]. Although SVs identification is a key step to grasp the “missing heritability” (not explained by SNPs) underlying complex traits involved in major evolutionary processes such as speciation, phenotypic plasticity, and adaptive responses, SV-based studies are too few mainly due to the lack of reliable detection methods [3]–[7]. The rise of the next-generation high-throughput DNA sequencing technologies opened up new possibilities to access these untapped genomic variations. However, despite a plentiful supply of SVs callers, high-quality SVs mapping from sequencing data remains a scavenger hunt. SV callers' difficulty in characterizing precisely SVs by inferring breakpoint positions is due to the inherent integrity of short-read mapping signals, including read-pair, split-read and read-depth, which rely on their algorithms [8]. Therefore, these tools suffer from low sensitivity (30–70%) and high false-positive rate (up to 85%), and even combinatorial approaches based on the union or the intersection of detected SVs will result in higher false positive and higher false-negative rates, respectively [7]. New methods are required to overcome these hurdles. Here we present a machine learning-based approach using a Random Forest (RF) method for accurate deletion identification: RF4Del. Our approach hinges on a different paradigm. Instead of basing SVs detection on one or a couple of alignment signals (read-pair, split-read, read-depth) as the vast majority of callers, RF4Del “learns” from 13 mapping features to identify a deletion-specific pattern used afterward to perform deletion detections.

## Results & Discussion

### Deletion detection approach

Our approach is composed of two main parts. The first part is dedicated to generating the prediction model using input mapping data. Once the prediction is done, the RF4Del tool can be used for the calling of deletions (**see Fig.1**). The prediction part starts by building the training matrix. To build the training matrix, twelve features were extracted from the simulated bam files from 12.7X sequencing data. The final training matrix was obtained by adding the variation event (deletion or not) for each position.

**Figure 1.**
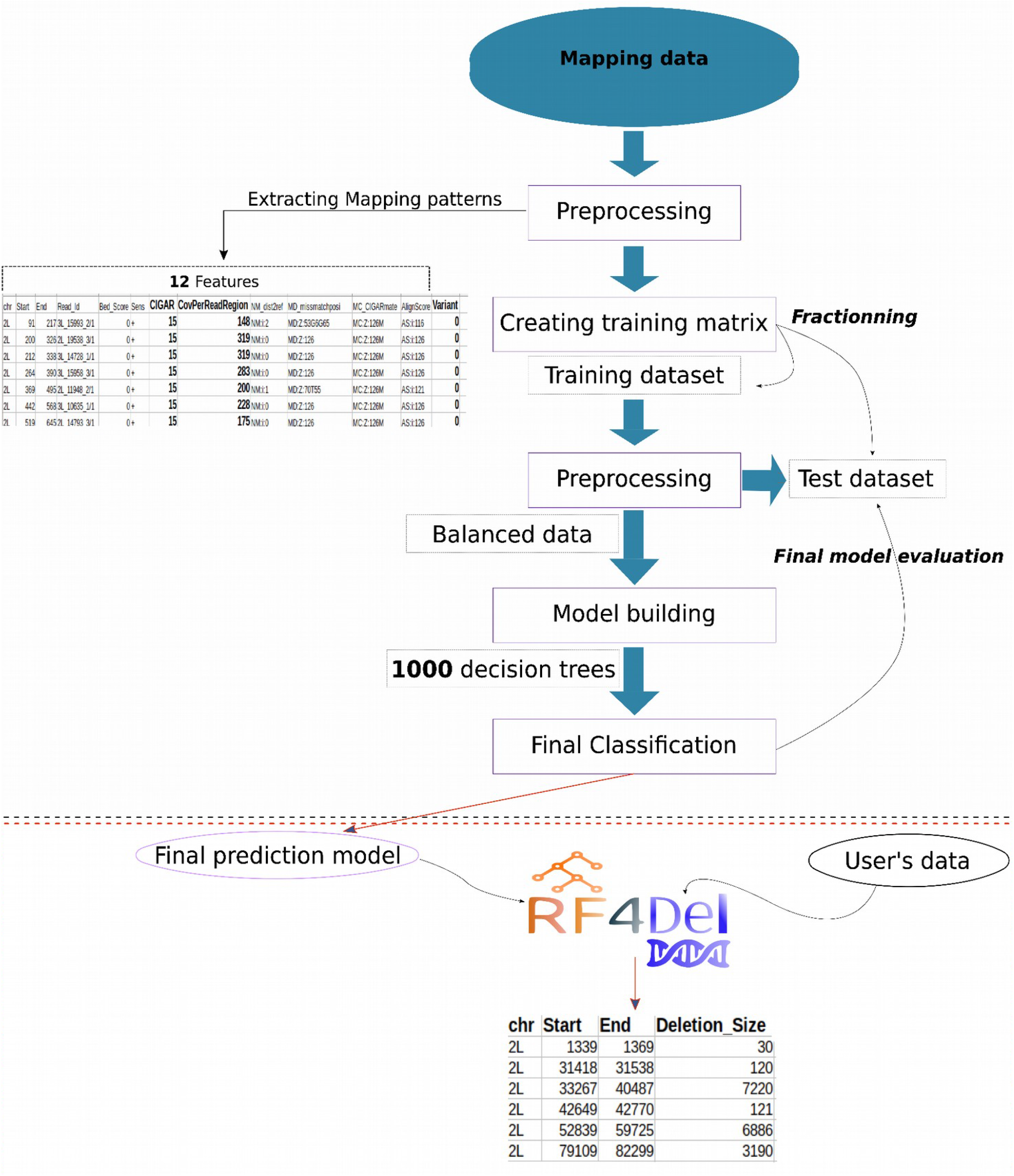
Two-step approach presented in this paper. The first step in the black dashed rectangle corresponds to the prediction model. Twelve mapping patterns are extracted from mapping data. The data are then fractionned to build 1000 decision trees and later the final classification. The second part in the red dashed rectangle corresponds to the deletion call by RF4Del using user’s data and based on the final prediction model.

To select the best method for the deletion detection, Machine Learning (ML) strategies, ranging from classic ones as Logistic Regression, Linear Discriminant Analysis, k-Nearest Neighbors, Naive Bayes Classification, and Regression Tree algorithm; to ensemble learning ones such as Adaptive Boosting classifier (AdaBoost), Gradient Boosting Machines (GBM) and Random Forest (RF) were evaluated (**see Supp Fig.1**). At the end of this first round of model testing, Linear Discriminant Analysis, Decision Tree, AdaBoost, RF and GBM reached the best results in all three metrics (**see Supp Fig.2**). Thus, to perform the subsequent testing, the three ensemble models (AdaBoost, RF, and GBM) were selected mainly because of their robustness when running with complex classification datasets. To compare the ensemble models against each other, 60% of the balanced dataset was used for training and 40% for testing. Running time was also added to the evaluation metrics. All three models reached almost 100% in all metrics, RF being the best model when considering the execution time **(see Supp Table1)**. To assess the impact of deep learning compared to neural learning, the RF model was then tested against neural networks with the same balanced dataset and with F1 score, Recall, and Precision as evaluation metrics. Stratified cross-validation (10 k-folds), and the train and test strategy were used to perform this final evaluation. Although the neural network models achieved high F1 score, Recall, and Precision scores, the RF model with equivalent scores outperformed in execution time **(see Supp Table1, Supp Table2)**. The deletion caller method has been devised on the basis of a RF model. We have explored the trade-off between model complexity and performance [9]. Therefore we have tried several RF models having distinct numbers of trees and features. No significant differences were detected between RF models configurations (**see Supp Fig.3**), most likely due to a balanced training dataset. The final prediction model built with 1000 decision trees is faster than a model with 100 decision trees (**see Supp Table3**). The training and building model steps took about 50 seconds (**see Supp Table4**).

### RF4Del can detect all deletions with accuracy

Testing SV callers using ground truth datasets is essential to assess a caller's performance and correctness. To test the veracity of our approach, we start by choosing a set of highly accurate validated deletions of different sizes. We used a set of 8 962 deletions detected in 39 lines derived from short-read DNA sequencing in a natural population (the “*Drosophila melanogaster* Genetic Reference Panel,” DGRP) [10]. And then we generated a simulated *D. melanogaster* genome using the reference genome (dm6). In order to be close to what we can expect in real nature populations, we launch the simulation taking into account the size of the 8 962 previously detected deletions from 50 bp to 165.32 kb length.

However, with a high prevalence of deletions smaller than 1 kb (37,6%: 3 376 out of 8 962) observed in the real data (**see Supp Fig.4.a**), we decided to simulate a set with larger deletions to assess accurately the deletion size effect on the detection. Comparison of the prevalence of deletions as a function of size between the “real” data from the study by Zichner *et al.* [10] and those from the simulation shows a size-biased prevalence in the real data and a more homogeneous prevalence in the real data and a more homogeneous prevalence in the simulated data (see Materials and Methods). We ended up with a simulated genome of 99,29 Mb with 8 962 deletions with the same length range but with a higher density of long deletions (**see Supp Fig.4.b**). We further simulated paired-end reads using this simulated genome with around 25X depth coverage that corresponds to approximately 20 million reads, comparable to what we obtained in the sequencing data from the natural *D. melanogaster* populations. The reads were then mapped on the reference genomic sequences (dm6; see Materials and Methods). 93% of the (8 309 out of the 8 962) deletions were correctly detected. The approach did not show a detection accuracy bias according to the deletion length.

### RF4Del can detect deletions with low coverage data

We further simulated paired-end reads using this simulated genome with lower depth coverage (1.2X, 6.3X and 12.7X, 25.3X). The short-reads from the *D. melanogaster* were then mapped on the reference genome (dm6). The final RF model was then loaded to predict new datasets with different sequencing coverage, 1.2X, 6.3X, 12.7X, and 25.4X corresponding to 1 million, 5 million, 10 million, and 20 million reads respectively (**see Supp Fig.5**). As expected the 1.2X sample returns the worst result with only 1 729 out of 8 962 deletions detected (19.3%). RF4Del shows a higher accuracy for the 6.3X (75.9%; 6 808 out of 8 962 deletions) and 12.7X (75.6%; 6 777 out of 8 962 deletions) samples than the 25X sample (67.8%; 6 078 out of 8 962 deletions). Previous studies confirm such results as they suggest how the increase of the data evidence biais the variation detection accuracy by adding noise [11]. A depth-analysis of the mapping for deletions detected using the 6.3X or 12.7X samples reveals putative deletions supported by numerous mapping evidence, not detected using the 25X sample. Even if the RF4Del sensibility for all the samples does increase with average coverages from 1.2X to 6.3X, the precision does not vary with the coverage and stay close to 100% (**see Supp Fig.6**). By consequence, 6.3X depth coverage data seems to be sufficient to accurately detect most of the deletions with RF4Del.

### RF4Del outperforms established SV callers

We then test the RF4Del veracity by comparing the detection results with those from well-known and commonly used variant callers: Pindel [12] and DELLY [13]. In contrast with RF4Del, the amount of detected deletions using DELLY and Pindel appears to be correlated with the amount of sequencing data. While DELLY detected around three-quarters of the deletions (73%; 6 542 out of 8 962), Pindel detected 54 190 deletions, suggesting a high number of false positives (**see Fig.2A**). With 6.3X coverage data, the amount of deletions detected using Pindel is more than twice the amount of expected deletions (**see Fig.2A**). The F1-score estimated for each dataset and tool confirms the low detection accuracy of Pindel. This analysis also highlights the higher sensitivity and precision of RF4Del with low coverage data compared with DELLY (**see Fig.2B; see Supp Fig.6**). Taken together, RF4Del appears as the best approach to detect most of the deletions with accuracy. A depth analysis allows estimating the F1 score using the 25.4X dataset using each tool per deletion size. The accuracy for the three approaches slightly decreases with the deletion length. Pindel fails to detect very short deletions (<100bp) and deletions from 500kb to 1.5kb length fail to be correctly detected by both DELLY and Pindel (**see Fig.2C; see Supp Fig.7**). The manual curation of the mapping in the vicinity of deletions of around 1kb reveals some mapping issues due to the insert size of the paired-end data.

**Figure 2.**
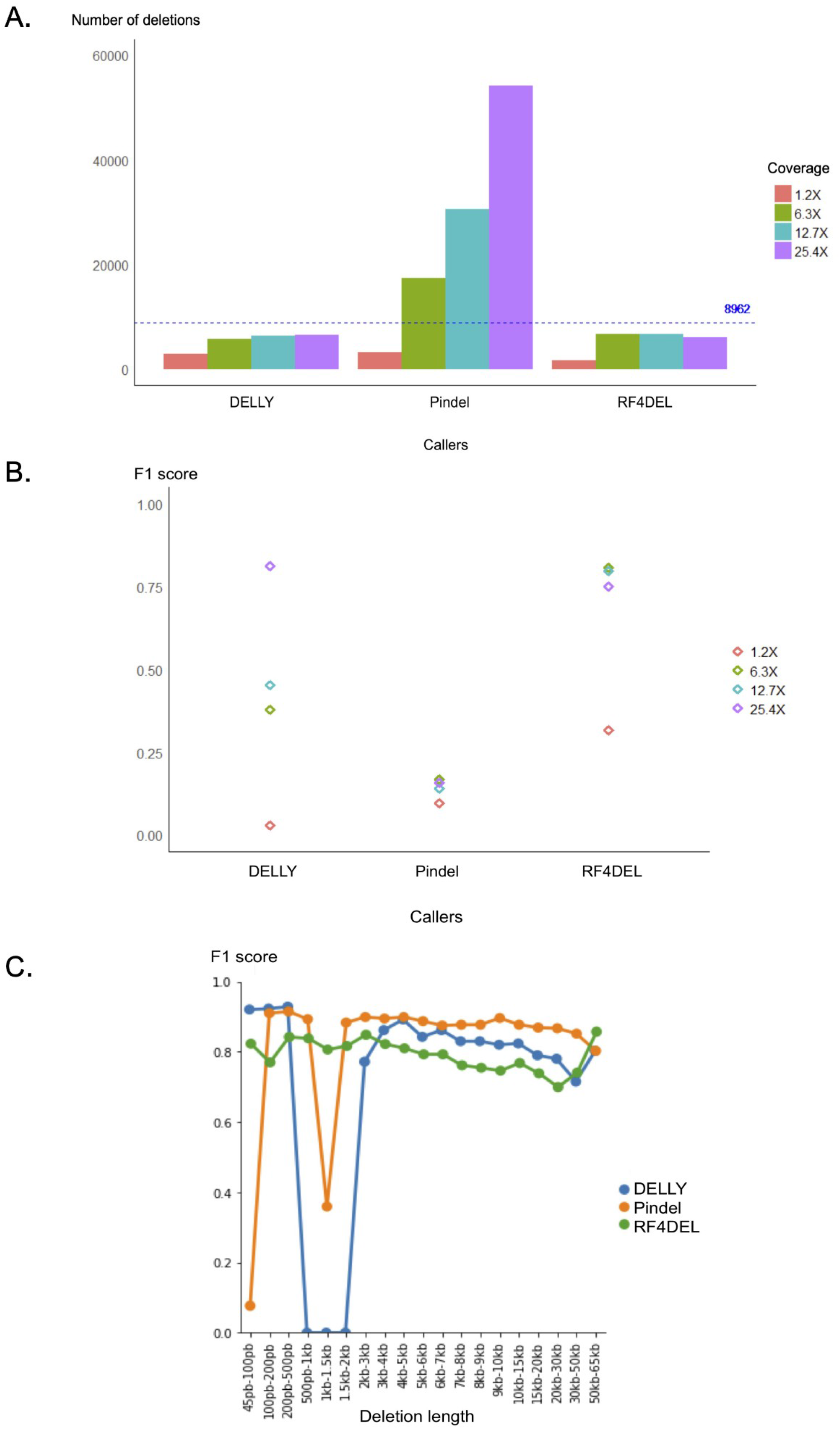
Comparison of the detection of deletions using different dataset and different tools. **A**. Number of deletions detected using the callers DELLY, Pindel and RF4Del with four datasets with variable coverage (1.2X, 6.3X, 12.7X, 25.4X). **B**. F1 score estimates based on all the deletions detected using the callers DELLY, Pindel and RF4Del with four datasets with variable coverage (1.2X, 6.3X, 12.7X, 25.4X). **C**. F1 score based on all the deletions detected using the 12.7X coverage dataset with the callers DELLY, Pindel and RF4Del per deletion length.

### Compared to other ML SV-callers, RF4Del is a good compromise for accurate deletion detection

Other similar tools which are also using a deep learning-based approach as RF4Del were recently developed: DeepSVFilter [14] and AquilaDeepFilter [15]. Also, a similar method was published in the preprint bioRxiv, called Scotch [16].

DeepSVFilter’s authors propose a deep learning-based approach for filtering SVs in short-reads sequencing data. In the training step, they use some state-of-the-art SV detection approaches, such as DELLY [13], Lumpy [17], and Manta [18], to detect SVs from data samples. Then DeepSVFilter encodes all SVs as images and trains CNN models. In the prediction module, they use the same strategy as the training module to generate SVs candidates and run CNN models to predict each candidate using a score between 0 and 1, representing the probability of the classification as the true SV. They adopt transfer learning with five state-of-the-art pre-trained CNN models on the ImageNet dataset [19] for training and prediction, including Inception-ResNet v2 [20], MobileNet v1 [21], MobileNet v2 [22], NASNet-A Mobile [23] and PNASNet-5 Mobile [24]*. However, this may not be the best approach as it takes considerable time in training and prediction steps with generated images. PNASNet-5 Mobile took about 4.87 hours and Inception-ResNet v2 took about 42.59 hours of training. Also, they run all CNN models on the server with the specifications: Intel(R) Xeon(R) Gold 5220R CPU 2.20GHz, 24 cores and 284 GB RAM, while RF4Del can run easily in a personal computer with AMD Ryzen 5 3500U 2.10GHz and 8GB RAM in few minutes (**see Supp Table1**). To demonstrate the classification accuracy of DeepSVFilter [14], they trained the model with 90% split of the dataset and used another 10% split to validate the trained model. In RF4Del, we trained our model with 60% split of the dataset and used 40% split to predict and validate the trained model, and reached almost 100% in F1 Score, Recall and Precision metrics.

AquilaDeepFilter filters large deletions from Aquila [25] and Aquila_stLFR [26]. Both tools are reference-assisted local assembly approaches to generate high-quality diploid assemblies and enable the genome-wide discovery and phasing of all types of variants including SNPs, small indels and SVs. However, they produce a substantial proportion of false positives, especially for large deletions. In this way, AquilaDeepFilter encodes the candidate variants from Aquila in RGB images splitted into two classes, candidate variants that are true deletions and candidate variants that are false-positive deletions, that will be the input of CNN Models. Then CNN models are trained in a supervised way and the prediction result of the model is stored in a BED file. In the evaluation step, they split the dataset in 90% for training and 10% for evaluation, and for all the experiments, a server with Intel(R) Xeon(R) CPU E5-2420 0 @ 1.90GHz, 24 cores and two high-performance GPU RTX 2080 Ti was used. Using this server, the training step took about 7 hours until convergence. In the results section, it is possible to see that AquilaDeepFilter generally improves the performance of Aquila and Aquila_stLFR by 20% in F1 Score metric, and also is compared with the previous approach DeepSVFilter. Using the same 10x lib and stLFR lib datasets, AquilaDeepFilter outperforms DeepSVFilter in most experiments in Precision, Recall and F1 Score metrics, however, there are some optimizations to increase general performance [14][15].

DeepSVFilter and AquilaDeepFilter show that Deep Learning using state-of-the-art CNNs can be applied to solve the problem of detecting SVs and InDels, respectively. Even nowadays we can run neural networks on our personal computer, complex applications still need a server with high-level configuration to do the experiments properly. In this case, the CNN models used took a high computational cost, and both approaches performed the experiments in two huge servers. One of the main advantages of RF4Del using RF is that it can solve this problem, more specifically in deletions, using a simple personal computer with AMD Ryzen 5 3500U 2.10GHz and 8 GB RAM. Another important aspect that must be considered using CNNs is the preprocessing of the images that DeepSVFilter and AquilaDeepFilter use as input in training and prediction steps. The preprocess is a step where important features of the data are encoded as RGB images, then a deep neural network model can be trained with this information. Training a model until it converges for good results so it can generalize and predict properly SVs and InDels in new data is a complex task that may take considerable running time. DeepSVFilter took about 4.87 hours with a simpler CNN and 42.59 hours with a more complex CNN. In its turn, AquilaDeepFilter took about 7 hours until convergence. In the RF4Del approach, images are unnecessary as it uses the important features of the data in a tabular file to train the model, consequently, the execution time is much faster, achieving convergence in only a few minutes. Nevertheless, we also tried a simple neural network model to compare with the main model. In Supplementary Table 2, we can see the results of the neural network experiment measured using stratified cross-validation with 10 k-folds for Recall, Precision and F1 Score, and the runtime was just over two minutes for each epoch. A second neural network experiment used 60% split data to train and 40% split data to test, and the model performed an average of: F1 score 0.9295%; Recall 0.9185%; and Precision 0.9534 in running time average of 145 seconds. RF4Del model’s average prediction time varies between 22 sec and 6 min 43 sec, which is much faster than the other tools, with a comparable amount of data.

Scotch is another ML-approach designed for the indel breakpoint detection, more similar to RF4Del as it also uses RF in its approach. In the pre-processing step, the tool examines designated sequence alignments (*i.e.*, mapping) and analyzes each base individually. Then, it creates a numerical profile of these positions, describing features of the aligned sequencing data like depth, base quality, and alignment to the reference. For the prediction, each position is classified based on the RF model as a non-indel or a type of indel breakpoint. The Scotch efficiency was evaluated with five other tools, such as DeepVariant [27], GATK [28], VarScan2 [29] and two versions of Pindel using three benchmark datasets: simulated variants, Syndip [30], and NA12878 [31]. For the deletion detection, Scotch outperformed both versions of Pindel with simulated data. However, its accuracy is quite low, as Scotch reaches only 38.2% of Precision and about 50% for F1 Metric [16]. In the supplementary section, analyzing the hyperparameters employed, authors considered four possible values for ntree (350, 450, 550, 650), and two for mtry (7, 8), resulting in eight possible hyperparameter configurations. In the RF4Del model, we considered the values 100, 200, 400, 800 and 1000 for ntree and 2, 4, 6, 8 and 10 for mtry, totalizing twenty-five different hyperparameter configurations. In Supplementary Figure 3, our experiments show that the model achieves 99.8% on F1 Score metric for each possible configuration. In addition, Supplementary Table 4 compares the running time of the RF4Del model with each ntree and mtry configuration, concluding that the faster took only 47 seconds with 100 ntree and 10 mtry. Scotch’s runtime is approximately 24 hours when parallelized by chromosomes with their adopted data. In the results, even Scotch’s performance in deletions with simulated data is better than both versions of Pindel, the accuracy is not satisfactory, as it reaches only 50% for F1 Score Metric. Comparing all these tools, RF4Del appears thus as the most appropriate tool to investigate with accuracy the detection of a short and large deletion in a fast and easy way with a personal computer (**see Table1**).

**Table 1.**
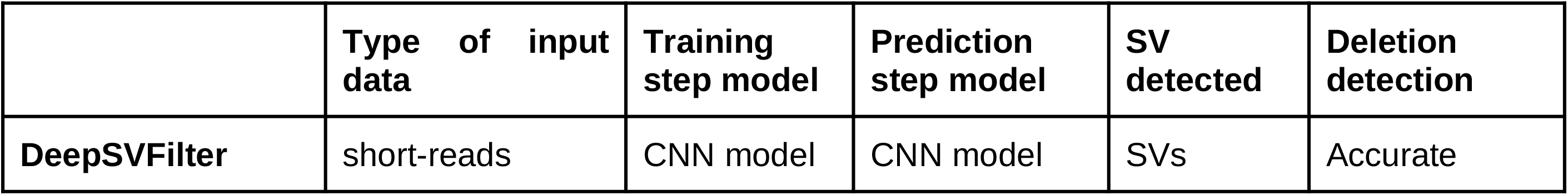
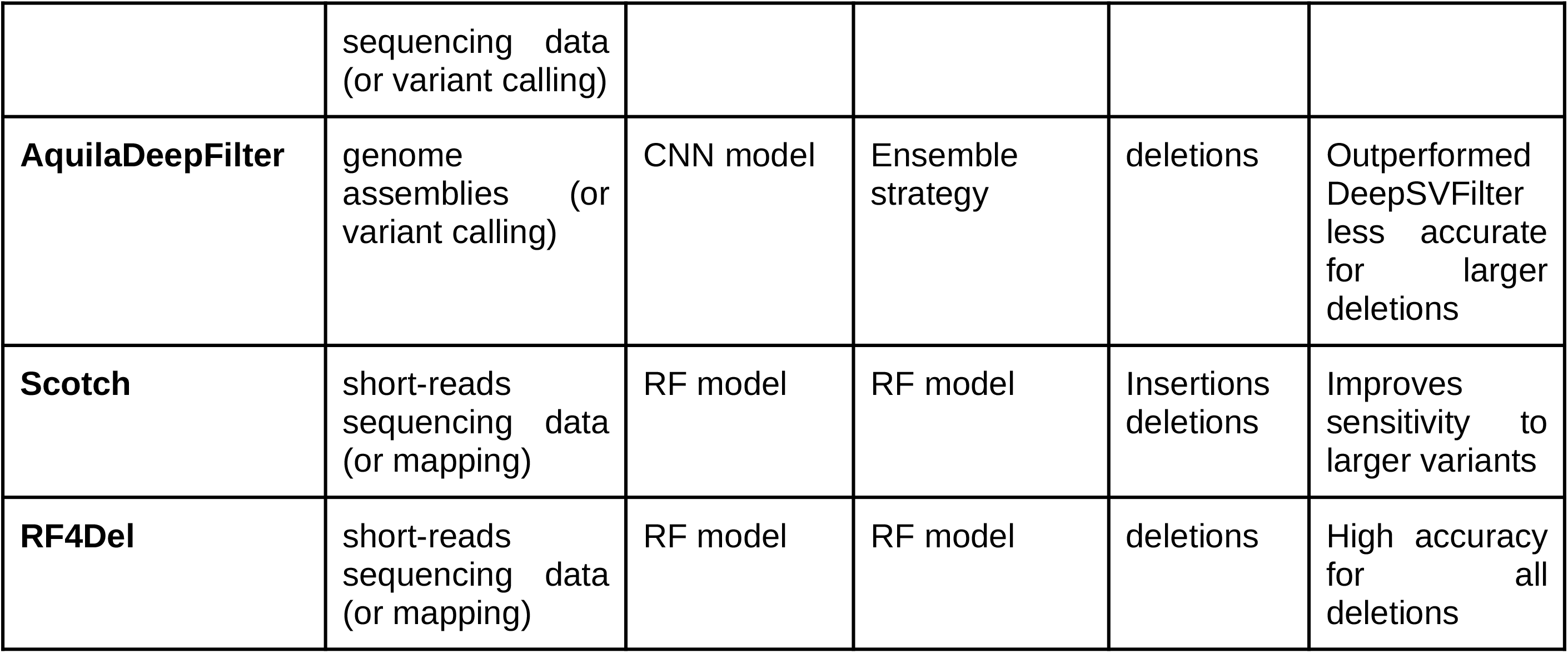
Comparison with recent other ML-based approach tools: DeepSVFilter [14], AquilaDeepFilter [15] and Scotch [16].

|  | Type of input data | Training step model | Prediction step model | SV detected | Deletion detection |
| --- | --- | --- | --- | --- | --- |
| DeepSVFilter | short-reads sequencing data<br>(or variant calling) | CNN model | CNN model | SVs | Accurate |
| **AquilaDeepFilter** | genome assemblies (or<br>variant calling) | CNN model | Ensemble<br>strategy | deletions | Outperformed<br>DeepSVFilter<br>less accurate<br>for larger<br>deletions |
| **Scotch** | short-reads<br>sequencing data<br>(or mapping) | RF model | RF model | Insertions<br>deletions | Improves<br>sensitivity to<br>larger variants |
| **RF4Del** | short-reads<br>sequencing data<br>(or mapping) | RF model | RF model | deletions | High accuracy<br>for all<br>deletions |

## Materials & Methods

### Genomics data

The *D. melanogaster* genome release 6 were downloaded (137.547Mb) on the ftp server: ftp://ftp.ensemblgenomes.org/pub/metazoa/release-43/fasta/drosophila_melanogaster/dna/Drosophila_melanogaster.BDGP6.22.dna.chromosome.*. The structural variations (SVs) catalog come from the study of Zichner *et al.* [10]. In this study, the authors used 39 *D. melanogaster* lines from the DGRP pilot data [32] (Illumina^®^ technology) to call SVs. The SVs detected by different tools such as DELLY and Pindel were then merged. Then some random SVs were experimentally validated. In the SV catalog, they detected 8 962 deletions from 50 pb to 165.32 kb (median 178 bp; **see Supp Fig. 4**).

### Simulated data

We simulated sequencing data from the *D. melanogaster* genome. Deletions were simulated using the *simulateSV* function of the *RSVsim* package (version 1.34)[33]. The size range and the number of simulated deletions were parameterized according to the SVs catalog. The *estimationSV* function of the RSVsim package (version 1.34) was used to draw 8 962 random deletions, sizes within the range from 50 pb to 61 kb according to a beta distribution and homogeneously distributed along the genome. The deletions are thus cut from the rearranged genome and their uniform breakpoints placed. The *inSilicoSeq* software [34] was used to obtain simulated reads from the rearranged genome (99.285 Mb). The Hiseq (Illumina®) error model was used to generate read-pairs of 126 bp and four sequencing coverage (1.2X, 6.3X, 12.7X, and 25.4X). These reads were then aligned to the *D. melanogaster* reference genome (not rearranged) using the default parameters of BWA-mem alignment software [35].

### Preparing training data set

To build the training matrix, 12 features were extracted from the simulated bam files from 12.7X sequencing data (**see Supp tab 1**). Bam files were converted to bed files using BEDTools (version 2.30.0)[36] suite sub-command *BamtoBed* with −cigare option to extract the CIGAR information for each mapped read. *SAMtools bedcov* (version 1.13) [37] command was used to estimate the coverage for each mapped read position from the bed file. The remaining features were extracted from the bam files using *SAMtools view* command (version 1.13)[37]. The final training matrix was obtained by adding the variation event (deletion or not) for each position. The final training data set has 13 features (**see Supp tab 1**) along with 10 015 384 instances. The 13^th^ feature corresponds to the class label. Labels are defined according to a dummy transformation for the class either, as ‘No Deletion’ or ‘Deletion’. The data set mirrors biological reality, hence the predominant label being ‘No Deletion’ with 9 835 103 instances and the Deletion label representing 0.5% of all instances with 180 181. To handle such an imbalanced data set, data corresponding to ‘Deletion’ and ‘No Deletion’ was first separated from the original dataset, then random samples from each group were selected. A final balanced data set summing up a total of 700 000 instances (150.000 ‘Deletion’ (21.5%) and 550.000 ‘No Deletion’ (78.5%)) was obtained. The CIGAR and label values were encoded using an *one_hot_encoding* strategy transforming each categorical value into a numerical one.

### Benchmarking

The benchmark of the ML methods was performed by the machine learning-dedicated Python library, *scikit-learn* (version 0.22) [38] using default parameters. RF4Del performances were benchmarked against two conventional callers relying on short-read mapping signals, DELLY (version 0.8.1) (RP and SR) [13] and Pindel (SR) [12] using F1 score, Recall, and Precision as metrics.

### The RF deletion prediction model

The final prediction model was built with 1000 decision trees. The model was saved in a. sav file that can be loaded anytime to perform prediction with new data.

## Conclusion & Perspectives

With new sequencing technologies advances, the study of the impact of structural variations in genome evolution is more and more affordable. Deletions represent a large part of structural variants. They can carry genes and by consequence change the gene expression or composition. Despite their effect on phenotype was not well studied. Approaches to detect genomics deletion might fail for some deletions based on their length and the amount of supporting evidence [39]. By consequence, current genome-wide deletion detection combines several tools that induce a large effort in terms of installing but also running time. Also as some organisms reveal preferentially large deletions with a specific evolutionary history, one might need to select the best tools to combine. The *in vitro* experimentations generating large real datasets with specific features are tedious and cost-prohibitive [31]. In silico simulations, on the other hand, are a low-cost, unbiased alternative and unlimited generating data allowing accurate Precision and Recall estimations of SV calling methods.

Here, we design a new learning-based approach to detect all types of deletions that we call RF4Del. This tool is based on an RF method for accurate deletion identification starting with mapping data. Taking all together, RF4Del overcomes the main deletion detection limits as it can be used to call deletions using short-read data, with very low coverage in a fast way (**see Table1**). With the exponential increase of sequencing projects for which we are interested in the impact of deletions on evolutionary processes, we need tools to detect the SVs with accuracy and in a reasonable amount of time. As the features from the long-read mapping differ, for the moment, RF4Del cannot be used to detect deletions using long-read sequencing data. Thus, the next step will be to see how to adapt RF4Del or use the same approach to call other types of SVs using all types of sequencing data.

## Supporting information

see Supplementary information

## Acknowledgements

Part of this work was supported by “La Région Occinatie”.

## References

[1] M. J. P. Chaisson et al., « Multi-platform discovery of haplotype-resolved structural variation in human genomes », Nat. Commun., vol. 10, n° 1, p. 1784, déc. 2019, doi: 10.1038/s41467-018-08148-z.

[2] P. D. Stenson et al., « The Human Gene Mutation Database: towards a comprehensive repository of inherited mutation data for medical research, genetic diagnosis and next-generation sequencing studies », Hum. Genet., vol. 136, n° 6, p. 665–677, juin 2017, doi: 10.1007/s00439-017-1779-6.

[3] T. A. Manolio et al., « Finding the missing heritability of complex diseases », Nature, vol. 461, n° 7265, p. 747–753, oct. 2009, doi: 10.1038/nature08494.

[4] E. E. Eichler et al., « Missing heritability and strategies for finding the underlying causes of complex disease », Nat. Rev. Genet., vol. 11, n° 6, p. 446–450, juin 2010, doi: 10.1038/nrg2809.

[5] J. L. Bennetzen et H. Wang, « The Contributions of Transposable Elements to the Structure, Function, and Evolution of Plant Genomes », Annu. Rev. Plant Biol., vol. 65, n° 1, p. 505–530, avr. 2014, doi: 10.1146/annurev-arplant-050213-035811.

[6] F. Marroni, S. Pinosio, et M. Morgante, « Structural variation and genome complexity: is dispensable really dispensable? », Curr. Opin. Plant Biol., vol. 18, p. 31–36, avr. 2014, doi: 10.1016/j.pbi.2014.01.003.

[7] F. J. Sedlazeck, H. Lee, C. A. Darby, et M. C. Schatz, « Piercing the dark matter: bioinformatics of long-range sequencing and mapping », Nat. Rev. Genet., vol. 19, n° 6, p. 329–346, juin 2018, doi: 10.1038/s41576-018-0003-4.

[8] D. E. Larson et al., « svtools: population-scale analysis of structural variation », Bioinformatics, vol. 35, n° 22, p. 4782–4787, nov. 2019, doi: 10.1093/bioinformatics/btz492.

[9] Z. Khan et al., « Ensemble of optimal trees, random forest and random projection ensemble classification », Adv. Data Anal. Classif., vol. 14, n° 1, p. 97–116, mars 2020, doi: 10.1007/s11634-019-00364-9.

[10] T. Zichner et al., « Impact of genomic structural variation in *Drosophila melanogaster* based on population-scale sequencing », Genome Res., vol. 23, n° 3, p. 568–579, mars 2013, doi: 10.1101/gr.142646.112.

[11] S. Sandmann et al., « Evaluating Variant Calling Tools for Non-Matched Next-Generation Sequencing Data », Sci. Rep., vol. 7, n° 1, p. 43169, mars 2017, doi: 10.1038/srep43169.

[12] K. Ye, M. H. Schulz, Q. Long, R. Apweiler, et Z. Ning, « Pindel: a pattern growth approach to detect break points of large deletions and medium sized insertions from paired-end short reads », Bioinformatics, vol. 25, n° 21, p. 2865–2871, nov. 2009, doi: 10.1093/bioinformatics/btp394.

[13] T. Rausch, T. Zichner, A. Schlattl, A. M. Stutz, V. Benes, et J. O. Korbel, « DELLY: structural variant discovery by integrated paired-end and split-read analysis », Bioinformatics, vol. 28, n° 18, p. i333–i339, sept. 2012, doi: 10.1093/bioinformatics/bts378.

[14] Y. Liu, Y. Huang, G. Wang, et Y. Wang, « A deep learning approach for filtering structural variants in short read sequencing data », Brief. Bioinform., vol. 22, n° 4, p. bbaa370, juill. 2021, doi: 10.1093/bib/bbaa370.

[15] Y. Hu, S. V. Mangal, L. Zhang, et X. Zhou, « An ensemble deep learning framework to refine large deletions in linked-reads », Genomics, preprint, sept. 2021. doi: 10.1101/2021.09.27.462057.

[16] C. Curnin et al., « Machine learning-based detection of insertions and deletions in the human genome », Bioinformatics, preprint, mai 2019. doi: 10.1101/628222.

[17] R. M. Layer, C. Chiang, A. R. Quinlan, et I. M. Hall, « LUMPY: a probabilistic framework for structural variant discovery », Genome Biol., vol. 15, n° 6, p. R84, 2014, doi: 10.1186/gb-2014-15-6-r84.

[18] X. Chen et al., « Manta: rapid detection of structural variants and indels for germline and cancer sequencing applications », Bioinformatics, vol. 32, n° 8, p. 1220–1222, avr. 2016, doi: 10.1093/bioinformatics/btv710.

[19] J. Deng, W. Dong, R. Socher, L.-J. Li, Kai Li, et Li Fei-Fei, « ImageNet: A large-scale hierarchical image database », in 2009 IEEE Conference on Computer Vision and Pattern Recognition, Miami, FL, juin 2009, p. 248–255. doi: 10.1109/CVPR.2009.5206848.

[20] C. Szegedy, S. Ioffe, V. Vanhoucke, et A. Alemi, « Inception-v4, Inception-ResNet and the Impact of Residual Connections on Learning », 2016, doi: 10.48550/ARXIV.1602.07261.

[21] A. G. Howard et al., « MobileNets: Efficient Convolutional Neural Networks for Mobile Vision Applications », ArXiv170404861 Cs, avr. 2017, Consulté le: 9 mars 2022. [En ligne]. Disponible sur: http://arxiv.org/abs/1704.04861

[22] M. Sandler, A. Howard, M. Zhu, A. Zhmoginov, et L.-C. Chen, « MobileNetV2: Inverted Residuals and Linear Bottlenecks », ArXiv180104381 Cs, mars 2019, Consulté le: 9 mars 2022. [En ligne]. Disponible sur: http://arxiv.org/abs/1801.04381

[23] B. Zoph, V. Vasudevan, J. Shlens, et Q. V. Le, « Learning Transferable Architectures for Scalable Image Recognition », ArXiv170707012 Cs Stat, avr. 2018, Consulté le: 10 mars 2022. [En ligne]. Disponible sur: http://arxiv.org/abs/1707.07012

[24] C. Liu et al., « Progressive Neural Architecture Search », ArXiv171200559 Cs Stat, juill. 2018, Consulté le: 10 mars 2022. [En ligne]. Disponible sur: http://arxiv.org/abs/1712.00559

[25] X. Zhou, L. Zhang, Z. Weng, D. L. Dill, et A. Sidow, « Aquila enables reference-assisted diploid personal genome assembly and comprehensive variant detection based on linked reads », Nat. Commun., vol. 12, n° 1, p. 1077, déc. 2021, doi: 10.1038/s41467-021-21395-x.

[26] Y. H. Liu et al., « Aquila_stLFR: diploid genome assembly based structural variant calling package for stLFR linked-reads », Bioinforma. Adv., vol. 1, n° 1, p. vbab007, juin 2021, doi: 10.1093/bioadv/vbab007.

[27] R. Poplin et al., « A universal SNP and small-indel variant caller using deep neural networks », Nat. Biotechnol., vol. 36, n° 10, p. 983–987, nov. 2018, doi: 10.1038/nbt.4235.

[28] A. McKenna et al., « The Genome Analysis Toolkit: A MapReduce framework for analyzing next-generation DNA sequencing data », Genome Res., vol. 20, n° 9, p. 1297–1303, sept. 2010, doi: 10.1101/gr.107524.110.

[29] D. C. Koboldt et al., « VarScan 2: Somatic mutation and copy number alteration discovery in cancer by exome sequencing », Genome Res., vol. 22, n° 3, p. 568–576, mars 2012, doi: 10.1101/gr.129684.111.

[30] H. Li et al., « A synthetic-diploid benchmark for accurate variant-calling evaluation », Nat. Methods, vol. 15, n° 8, p. 595–597, août 2018, doi: 10.1038/s41592-018-0054-7.

[31] J. M. Zook et al., « Integrating human sequence data sets provides a resource of benchmark SNP and indel genotype calls », Nat. Biotechnol., vol. 32, n° 3, p. 246–251, mars 2014, doi: 10.1038/nbt.2835.

[32] T. F. C. Mackay et al., « The Drosophila melanogaster Genetic Reference Panel », Nature, vol. 482, n° 7384, p. 173–178, févr. 2012, doi: 10.1038/nature10811.

[33] C. Bartenhagen et M. Dugas, « RSVSim: an R/Bioconductor package for the simulation of structural variations », Bioinformatics, vol. 29, n° 13, p. 1679–1681, juill. 2013, doi: 10.1093/bioinformatics/btt198.

[34] H. Gourlé, O. Karlsson-Lindsjö, J. Hayer, et E. Bongcam-Rudloff, « Simulating Illumina metagenomic data with InSilicoSeq », Bioinformatics, vol. 35, n° 3, p. 521–522, févr. 2019, doi: 10.1093/bioinformatics/bty630.

[35] H. Li et R. Durbin, « Fast and accurate short read alignment with Burrows-Wheeler transform », Bioinforma. Oxf. Engl., vol. 25, n° 14, p. 1754–1760, juill. 2009, doi: 10.1093/bioinformatics/btp324.

[36] A. R. Quinlan et I. M. Hall, « BEDTools: a flexible suite of utilities for comparing genomic features », Bioinformatics, vol. 26, n° 6, p. 841–842, mars 2010, doi: 10.1093/bioinformatics/btq033.

[37] H. Li et al., « The Sequence Alignment/Map format and SAMtools », Bioinformatics, vol. 25, n° 16, p. 2078–2079, août 2009, doi: 10.1093/bioinformatics/btp352.

[38] L. Buitinck et al., « API design for machine learning software: experiences from the scikit-learn project », ArXiv13090238 Cs, sept. 2013, Consulté le: 17 février 2022. [En ligne]. Disponible sur: http://arxiv.org/abs/1309.0238

[39] N. Wang et al., « Tool evaluation for the detection of variably sized indels from next generation whole genome and targeted sequencing data », PLOS Comput. Biol., vol. 18, n° 2, p. e1009269, févr. 2022, doi: 10.1371/journal.pcbi.1009269.

[40] W. Lin, Z. Wu, L. Lin, A. Wen, et J. Li, « An Ensemble Random Forest Algorithm for Insurance Big Data Analysis », IEEE Access, vol. 5, p. 16568–16575, 2017, doi: 10.1109/ACCESS.2017.2738069.

