## Supplementary material for "RF4Del: A Random Forest approach for accurate deletion detection": see Supplementary information

**Supplementary figures**

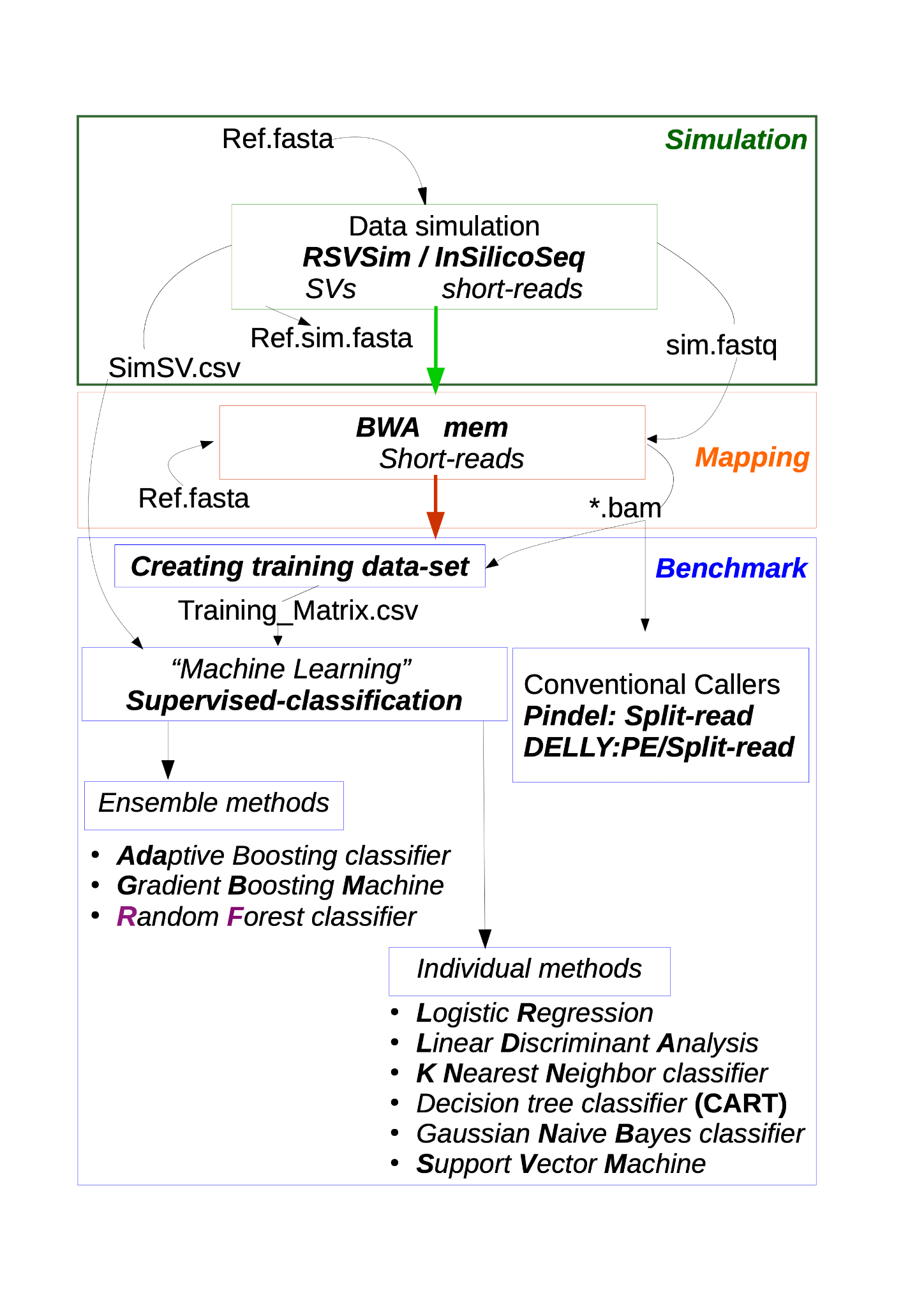

**Supp Figure 1.**
Approach used in the paper to select the best machine learning model. Using a reference sequence, structural variants (SVs) have been simulated. Then simulated short-reads have been generated from the new reference sequence. The short-reads were then mapped on the original reference sequence. Conventional callers were used to call the deletions. The mapping data were further used to create the training matrix. This matrix was finally used for a supervised classification based on ensemble and individual methods.

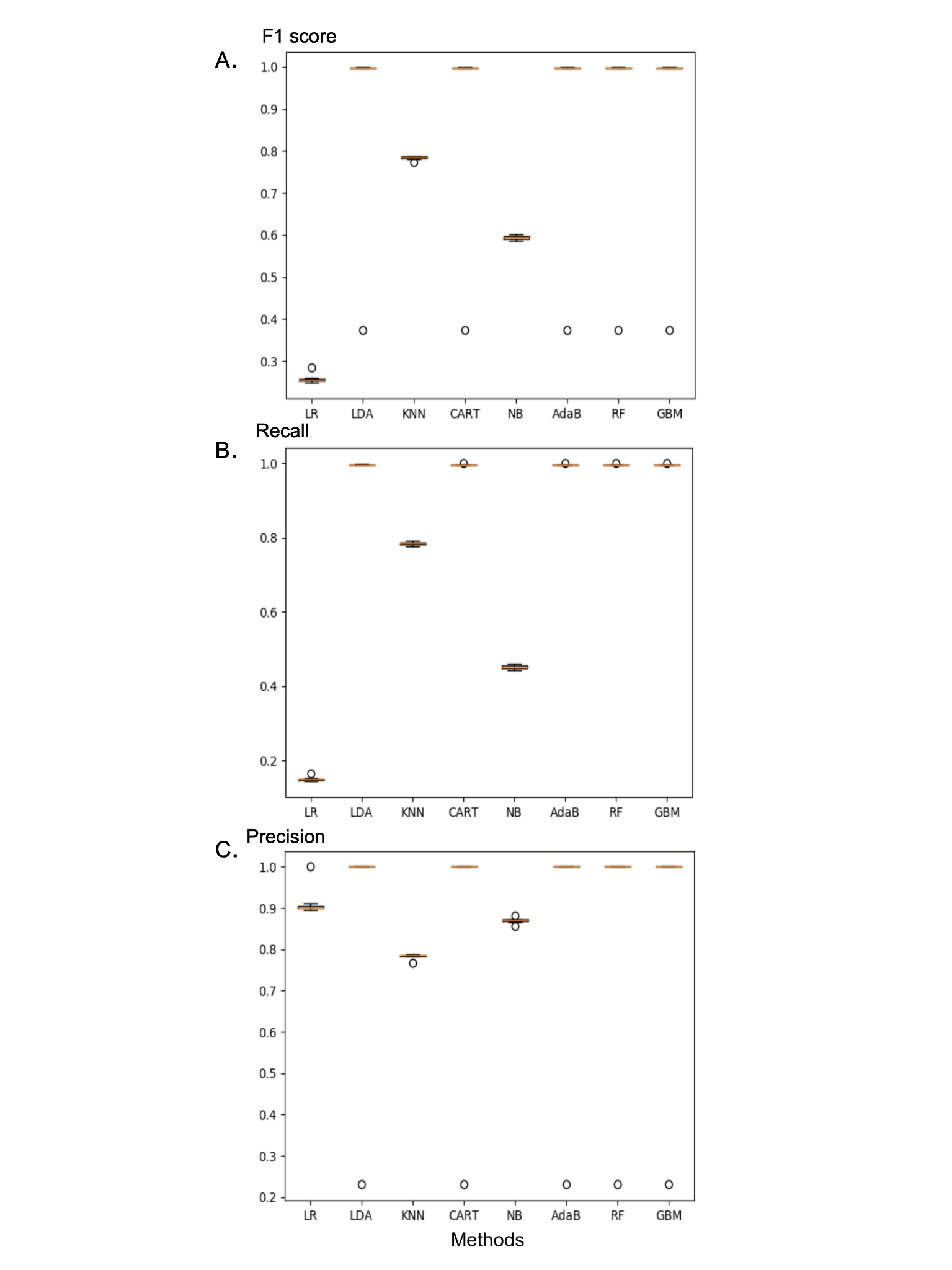

**Supp Figure 2.**

Performance comparison of models. All figures show boxplot graphs for the 8 models (LR : Logistic Regression; LDA: Linear Discriminant Analysis; KNN: K-Nearest Neighbor classifier; CART for Decision tree; NB: Naive Bayes classifier; AdaB : Adaptive Boosting classifier; RF : Random Forest classifier; GBM: Gradient Boosting Machine) **A** - Using F1 measure. The F1 score is an average of the precision and recall, where the F1 measure reaches its best value at 1 and worst value at 0. **B -** Using recall measure. The recall measure refers to the results correctly classified by the model. **C -** Using precision measure which indicates the percentage of the results which are relevant.

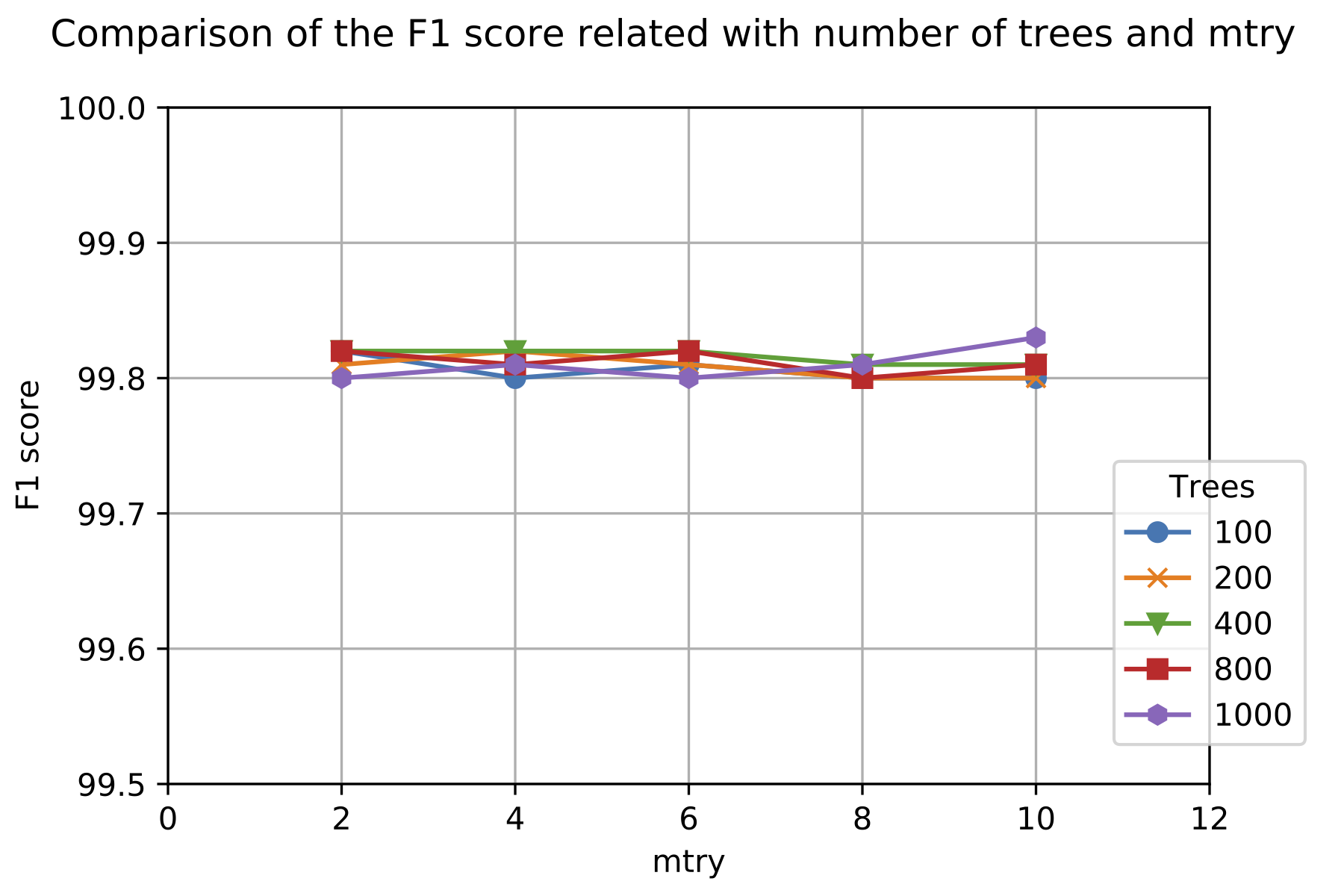
**Supp Figure 3.**
Evaluating RF models's complexity and performance. RF model with 1000 trees present a good compromise between complexity and performance (max number of features=mtry).

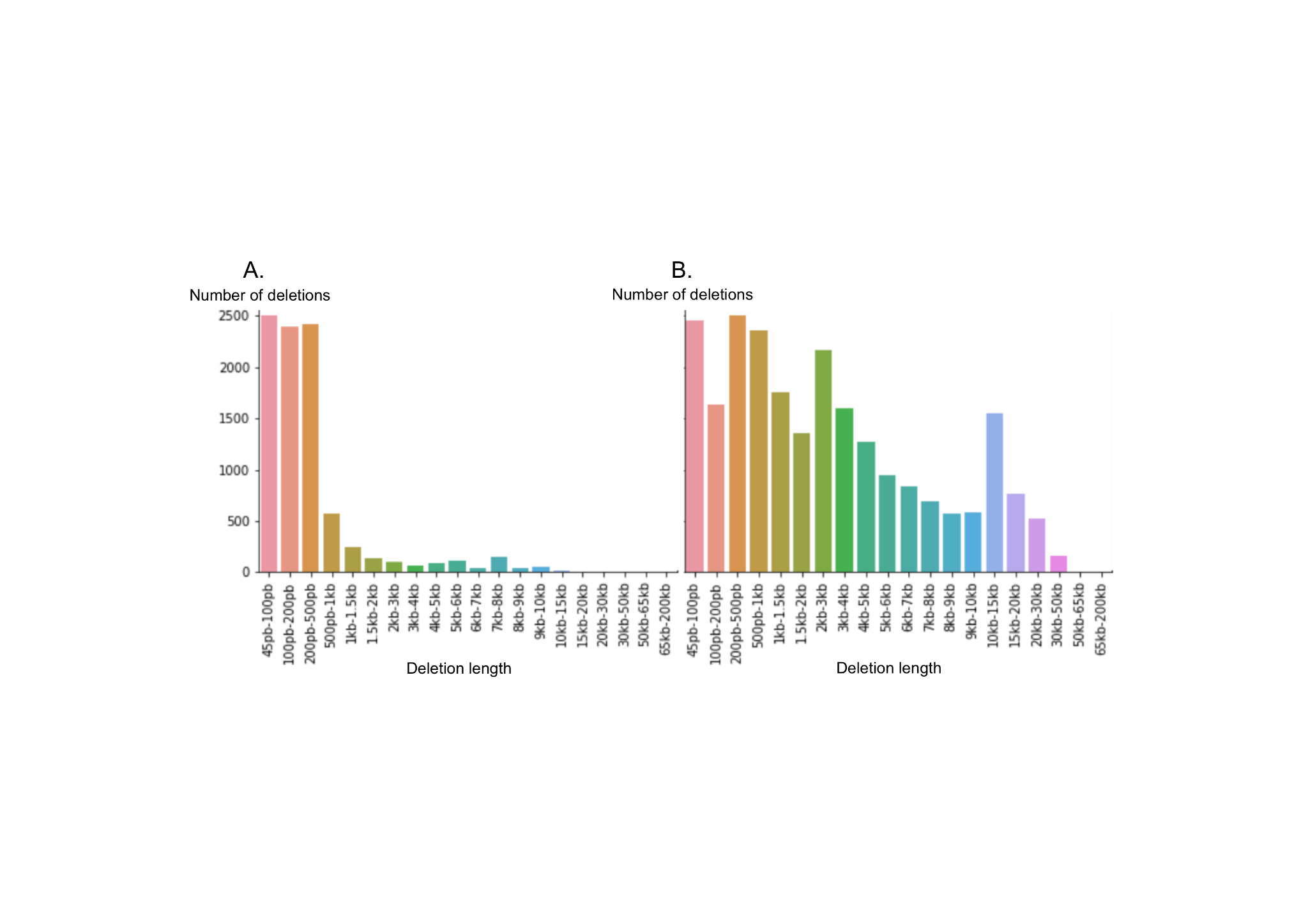

**Supp Figure 4.**

Distribution of the deletion length. **A -** real deletions. **B** - simulated deletions.

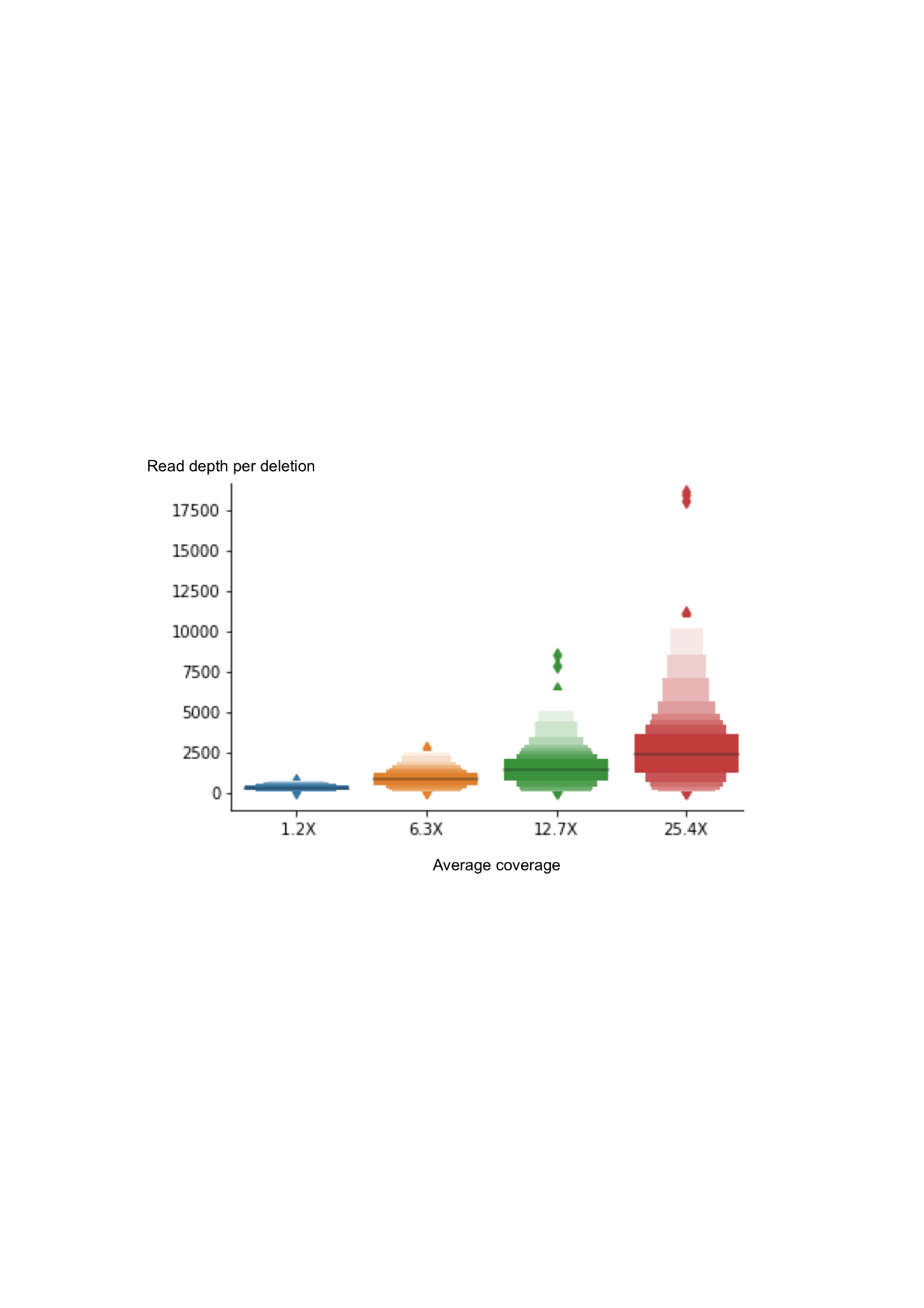

**Supp Figure 5**.

Read depth as a function of average coverage for the detected deletions. The read depths per deletion were calculated for four datasets that differ only in terms of the amount of reads: 1.2X, 6.3X, 12.7X, 25.4X.

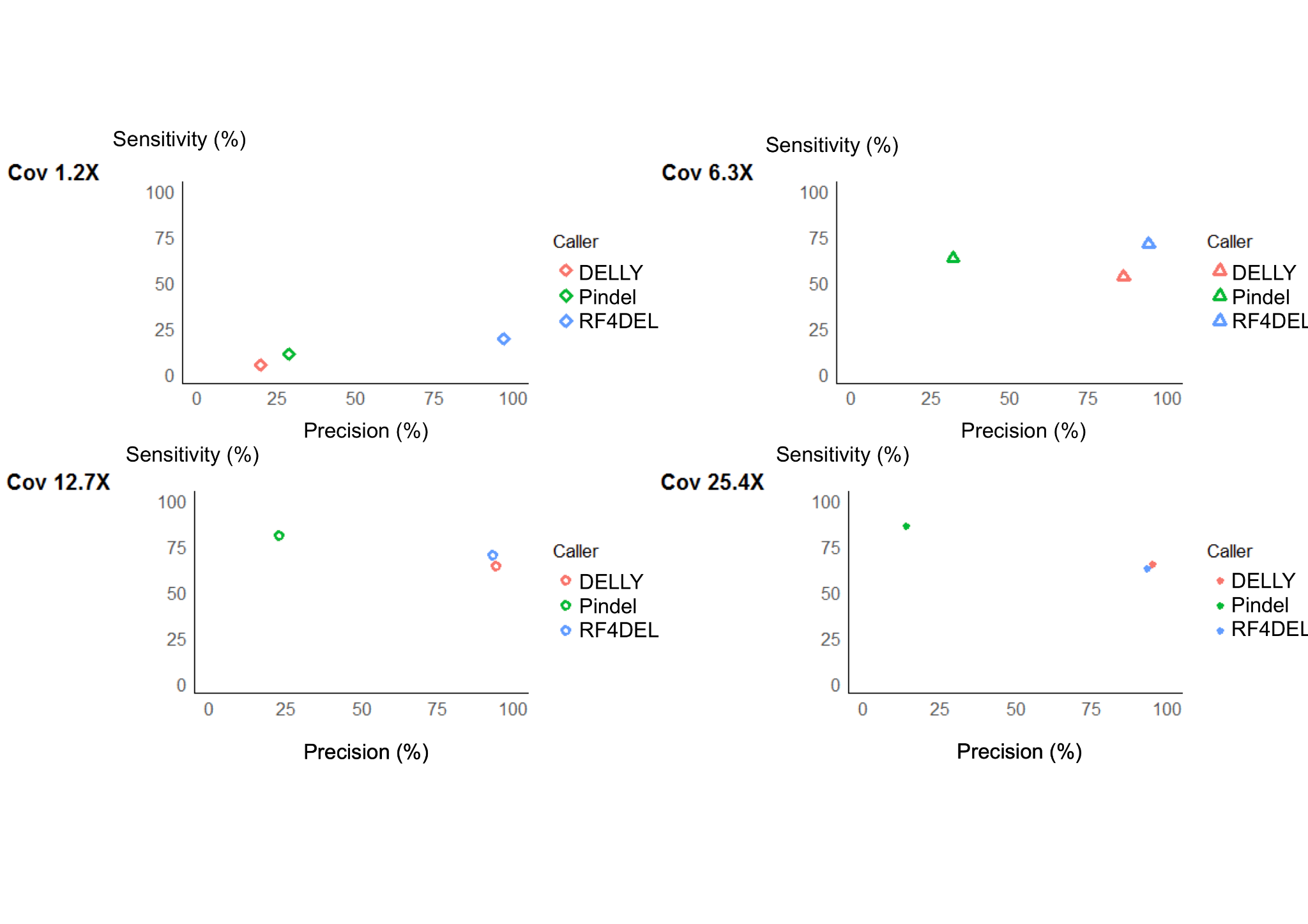

**Supp Figure 6.**

Sensitivity (%) as a function of the precision (%) for each variant callers (DELLY, Pindel and RF4Del) for each sample (1.2X, 6.3X, 12.7X and 25.4X).

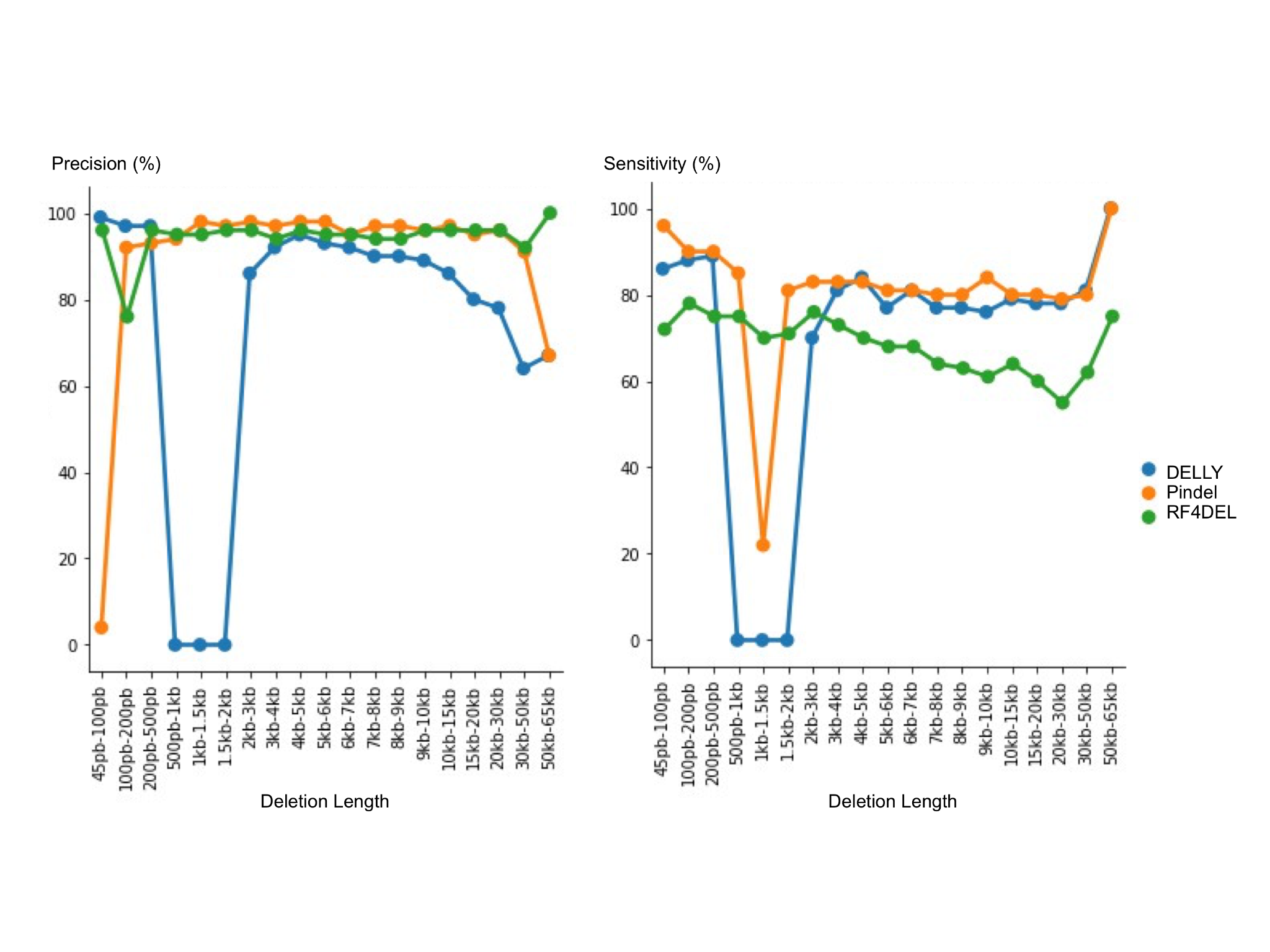

**Supp Figure 7.**

Precision and sensitivity for the deletions detected as a function of the deletion length (kb) for the three variant callers (DELLY, Pindel and RF4Del).

**Supplementary tables**

To build the training matrix, 12 features were extracted from the simulated bam files from 12.7X sequencing data (**see** **Supp Table 1**).

In this experiment, we use as input data the main balanced dataset in all three models with default parameters settings, and the strategie is to use 60% of the dataset to train and 40% to test. The experiments were performed using the default parameters of each model, it means, that the number of estimators was 100 trees for Random Forest (RF) model, 50 estimators for AdaBoost model, and 100 estimators for Gradient Boost model \cite{scikit-learn}. Table 1 below illustrates the results with F1 score, Recall, and Precision. Also, the running time was compared with the models. In Suup Table 1, it is possible to see that all three models reached almost 100% in all metrics, but RF is the fastest considering the time of execution. Therefore, the final model is based on the RF algorithm.

**Supp Table 1 -** Models evaluation considering F1 score, recall, precision and running time.

|  | RF (n_estimators=100) | | | AdaBoost (n_estimators=50) | | | GBM (n_estimators=100) | | |
| --- | --- | --- | --- | --- | --- | --- | --- | --- | --- |
|  | Run 1 | Run 2 | Run 3 | Run 1 | Run 2 | Run 3 | Run 1 | Run 2 | Run 3 |
| F1 score | 0.998237 | 0.998359 | 0.998149 | 0.998071 | 0.998324 | 0.998273 | 0.998044 | 0.998171 | 0.998013 |
| Recall | 0.996647 | 0.996888 | 0.996454 | 0.996150 | 0.996704 | 0.996569 | 0.996228 | 0.996399 | 0.996100 |
| Precision | 0.999833 | 0.999833 | 0.999850 | 1.000000 | 0.999950 | 0.999983 | 0.999867 | 0.999950 | 0.999933 |
| Time (in seconds) | 12.9855 | 12.8441 | 12.8071 | 74.8001 | 60.4933 | 28.4554 | 142.4562 | 74.4302 | 142.2374 |

The neural networks were also tested in some experiments to compare the performance with the RF model. The experiments were conducted with the same balanced dataset and with F1 score, Recall and Precision metrics to evaluate the neural network based model. Table 2 illustrates the results of the first experiment that were measured using stratified cross validation with 10 k-folds for all three metrics. The second experiment used the train and test strategy instead of cross validation, it means that for this experiment we split the dataset in 60% to train and 40% to test. For this experiment the model presented an average of: F1 score 0.9295%; Recall 0.9185%; and Precision 0.9534 in running time average of 145 seconds. We can notice, based on the first experiment, that the metrics are close to the RF model, but it took more time to execute. In the second experiment we can see that the evaluation was not as good as the RF model.

**SuppTable 2 - Neural network evaluation with stratified cross validation**

|  | F1 score | Recall | Precision | Loss | Time  (in seconds) |
| --- | --- | --- | --- | --- | --- |
| Epoch 1 | 0.8574 | 0.8991 | 0.8381 | 0.1250 | 136 |
| Epoch 2 | 0.9499 | 0.9702 | 0.9381 | 0.0699 | 129 |
| Epoch 3 | 0.9866 | 0.9893 | 0.9859 | 0.0300 | 129 |
| Epoch 4 | 0.9889 | 0.9889 | 0.9907 | 0.0221 | 129 |
| Epoch 5 | 0.9901 | 0.9886 | 0.9932 | 0.0186 | 129 |
| Epoch 6 | 0.9907 | 0.9885 | 0.9943 | 0.0164 | 129 |
| Epoch 7 | 0.9913 | 0.9889 | 0.9951 | 0.0149 | 129 |
| Epoch 8 | 0.9916 | 0.9890 | 0.9956 | 0.0137 | 129 |
| Epoch 9 | 0.9919 | 0.9893 | 0.9959 | 0.0127 | 129 |
| Epoch 10 | 0.9922 | 0.9896 | 0.9960 | 0.0119 | 129 |

The RF method consists of a large number of decision trees that work as a set, where each tree in the random forest predicts a class result and the class with the most votes becomes the model prediction [[40]](https://www.zotero.org/google-docs/?TEvHc3). The scikit-learn is a library for machine learning in Python that provides a large number of parameterization [[38]](https://www.zotero.org/google-docs/?tC5MFn). This library was used to create the RF model with 100 trees, its default. Other experiments were performed using distinct RF models, having more than 100 trees (**Supp** **Figure3**). RF from *Scikit-lear*n version 0.22 uses as a default 100 trees and “auto” as the max number of features (mtry). **Supp** **Figure 3** shows the experiments in setting up these two parameters, with a number of trees between 100 and 1000, and mtry from 2 to 10 max features. From all these different configurations, one can notice that models having 100 trees and 2 max features are performant and less complex than 1000 trees and 10 max features, respectively.

**Supp Table 3.** Random Forest model with 100 trees is faster than with 1000 trees.

| **Analysis of runnin time and F1 Score of Random Forest models ranging from 100 to 1000 trees and mtry from 2 to 10** | | | |
| --- | --- | --- | --- |
| **Trees** | **Mtry** | **F1 Score** | **Time (seconds)** |
| **1000** | 10 | 99.83 | 455.98 |
|  | 8 | 99.81 | 526.40 |
|  | 6 | 99.8 | 490.18 |
|  | 4 | 99.81 | 480.97 |
|  | 2 | 99.8 | 404.39 |
| **800** | 10 | 99.81 | 432.73 |
|  | 8 | 99.8 | 370.63 |
|  | 6 | 99.82 | 335.69 |
|  | 4 | 99.81 | 360.67 |
|  | 2 | 99.82 | 383.51 |
| **400** | 10 | 99.81 | 217.21 |
|  | 8 | 99.81 | 188.45 |
|  | 6 | 99.82 | 170.55 |
|  | 4 | 99.82 | 170.61 |
|  | 2 | 99.82 | 207.77 |
| **200** | 10 | 99.8 | 107.90 |
|  | 8 | 99.8 | 104.98 |
|  | 6 | 99.81 | 93.55 |
|  | 4 | 99.82 | 107.06 |
|  | 2 | 99.81 | 86.71 |
| **100** | 10 | 99.8 | 47.87 |
|  | 8 | 99.8 | 60.46 |
|  | 6 | 99.81 | 62.21 |
|  | 4 | 99.8 | 69.36 |
|  | 2 | 99.82 | 72.12 |

In addition to adding the runtime, in the supp Table 3, it is possible to notice almost no difference in F1 score along all the experiments, the F1 score varies from 99.8 to 99.83. However, it is possible to see a huge difference in runtime, it increases as long as the number of trees also increases. The mtry parameter made no difference in F1 score nor runtime. Therefore, it is better to maintain the default configuration since they are less complex, results are as good as the more complex models and faster.

**Supp Table 4.** Average time needed to predict new data with RF4Del model

|  | Dataset 1.2X | Dataset 6.3X | Dataset 12.7X | Dataset 25.4X |
| --- | --- | --- | --- | --- |
| Time (in seconds) | 22.64 | 115.03 | 229.25 | 409.25 |
